# The fluorescence-activating and absorption-shifting tag (FAST) enables live-cell fluorescence imaging of *Methanococcus maripaludis*

**DOI:** 10.1101/2022.04.01.486799

**Authors:** Eric Hernandez, Kyle C Costa

**Author notes:** Correspondence: Kyle C. Costa, Room 140, Gortner labs, 1479 Gortner Ave, St. Paul, MN 55108.

## Abstract

Live-cell fluorescence imaging in methanogenic archaea has been limited due to the strictly anoxic conditions required for growth and issues with autofluorescence associated with electron carriers in central metabolism. Here, we show that the fluorescence-activating and absorption-shifting tag (FAST) when complexed with the fluorogenic ligand 4-hydroxy-3-methylbenzylidene-rhodanine (HMBR) overcomes these issues and displays robust fluorescence in *Methanococcus maripaludis*. We also describe a mechanism to visualize cells under anoxic conditions using a fluorescence microscope. Derivatives of FAST were successfully applied for protein abundance analysis, subcellular localization, and determination of protein-protein interactions. FAST fusions to both formate dehydrogenase (Fdh) and F_420_-reducing hydrogenase (Fru) displayed increased fluorescence in cells grown on formate containing medium, consistent with previous studies suggesting increased abundance of these proteins in the absence of H_2_. Additionally, FAST fusions to both Fru and the ATPase associated with the archaellum (FlaI) showed membrane localization in single cells observed using anoxic fluorescence microscopy. Finally, a split reporter translationally fused to the alpha and beta subunits of Fdh reconstituted a functionally fluorescent molecule *in vivo* via bimolecular fluorescence complementation. Together, these observations demonstrate the utility of FAST as a tool for studying members of the methanogenic archaea.

**Importance:** Methanogenic archaea are important members of anaerobic microbial communities where they catalyze essential reactions in the degradation of organic matter. Developing additional tools for studying the cell biology of these organisms is essential to understanding them at a mechanistic level. Here, we show that FAST, in combination with the fluorogenic ligand HMBR, can be used to monitor protein dynamics in live cells of *M. maripaludis*. Application of FAST holds promise for future studies focused on the metabolism and physiology of methanogenic archaea.

## Introduction

Methanogenic archaea (methanogens) are responsible for producing the majority of methane on Earth and are model organisms for studying cellular processes in the Archaea. Several organisms such as *Methanococcus maripaludis, Methanosarcina* spp., and *Methanothermobacter* spp. have been used extensively in genetic or biochemical studies to understand the physiology and metabolism of methanogens (1–3). However, robust tools for direct cell visualization via fluorescence microscopy to allow for protein quantification, analysis of protein-protein interactions, and spatiotemporal characterization of cellular proteins have been lacking. A strict requirement for growth under anoxic conditions, as well as background autofluorescence due to the presence of the oxidized electron carrier coenzyme F_420_ (excitation peak centered at 420 nm, emission peak centered at 480 nm (4)), have prevented the use of most fluorescent protein tags. Green fluorescent protein (GFP) derivatives such as EGFP and mCherry require oxygen for fluorophore maturation and fail to perform under anaerobic growth conditions (5). Alternative oxygen independent probes such as FMN binding fluorescent proteins (e.g. LOV-based fluorescent proteins) typically have weak fluorescence intensity and often exhibit emission around 450 nm, overlapping with excitation/emission spectra of F_420_ (6). Immunofluorescence strategies using antibody-reporter conjugates typically involve aerobic steps and/or permeabilization of cells using harsh detergents, which preclude live-cell imaging of anaerobes. Another consequence of aerobic preparation of methanogenic organisms is that it causes the rapid oxidation of coenzyme F_420_ which leads to increased autofluorescence.

Recently, an oxygen independent fluorescent protein reporter known as the fluorescence-activating and absorption-shifting tag (FAST) has been developed and adapted in select anaerobic organisms (7–9). FAST is an engineered variant of photoactive yellow protein with a molecular mass of 14 kDa (7). Like GFP, FAST can be translationally fused to the N- or C-terminus of a gene of interest. By itself, FAST does not exhibit fluorescence; however, upon the addition of a fluorogenic ligand (fluorogen), the two complex together producing a fluorescent product. Multiple fluorogens consisting of 4-hydroxybenzylidene rhodanine (HBR) derivatives exist with each having unique excitation and emission properties upon binding to FAST, and variants of FAST have been developed that specifically bind a subset of fluorogens allowing for two color live-cell imaging (10, 11). HBR derivatives are membrane permeable and bind reversibly and specifically to FAST (7).

There are several reasons FAST is a particularly attractive molecular tool for studying obligate anaerobes such as methanogens. Most importantly, fluorescence is oxygen independent, and membrane permeability of the ligand allows for live-cell imaging. FAST complexes are inherently bright and emit fluorescence across a wide spectrum, avoiding the limitations of reporters such as FMN binding proteins. FAST fluorescence is reversible and quantitative, allowing for direct measurement of relative protein abundance (7, 12). Additionally, splitFAST was recently developed to visualize protein-protein interactions through bimolecular fluorescence complementation (BiFC) (13). Finally, as a small, translationally fused tag, FAST can be used to observe protein localization in live cells.

In this study, we demonstrate successful use of the FAST toolkit in *M. maripaludis*, demonstrating a functional fluorescence-based system for microscopic imaging in a methanogen. We developed a platform for microscopy under anoxic conditions, allowing for visualization of live cells. Combining these advances, we observed robust and quantifiable fluorescence of differentially expressed proteins in cells grown with either H_2_ or formate as an electron donor, BiFC using the multisubunit formate dehydrogenase (Fdh), and two different examples of protein localization. This was accomplished using the original FAST protein, also referred to as FAST1, as well as a highly fluorescent, tandem variant. FAST-based fluorescence microscopy expands existing tools for studying the cell biology of *M. maripaludis* and should be broadly applicable to other methanogens with established protocols for heterologous protein expression.

## Results

### Expression of FAST1 in *M. maripaludis*

To test the functionality of FAST, a codon optimized FAST1 gene (14) was expressed in *M. maripaludis* on the replicating vector pLW40 under control of the *Methanococcus voltae* histone promoter (15). Under these conditions, heterologously expressed protein can reach up to 1% of total cellular protein (16). 4-hydroxy-3-methylbenzylidene-rhodanine (HMBR), a FAST fluorogen, was added to stationary phase cultures (OD_600_ = ∼0.9) to a final concentration of 10 μM (based on manufacture’s recommendation) before transfer to a 96 well dark plate for quantification on a microplate reader. HMBR fluorescence was measured at 540 nm (excitation with 481 nm). Cultures that were not treated with HMBR exhibited little autofluorescence while cells containing both FAST1 and HMBR exhibited a significant increase in fluorescence (Fig. 1A). Addition of HMBR to wild type cells did not lead to a significant increase in fluorescence compared to cultures expressing FAST1 without HMBR addition.

**Figure 1.**
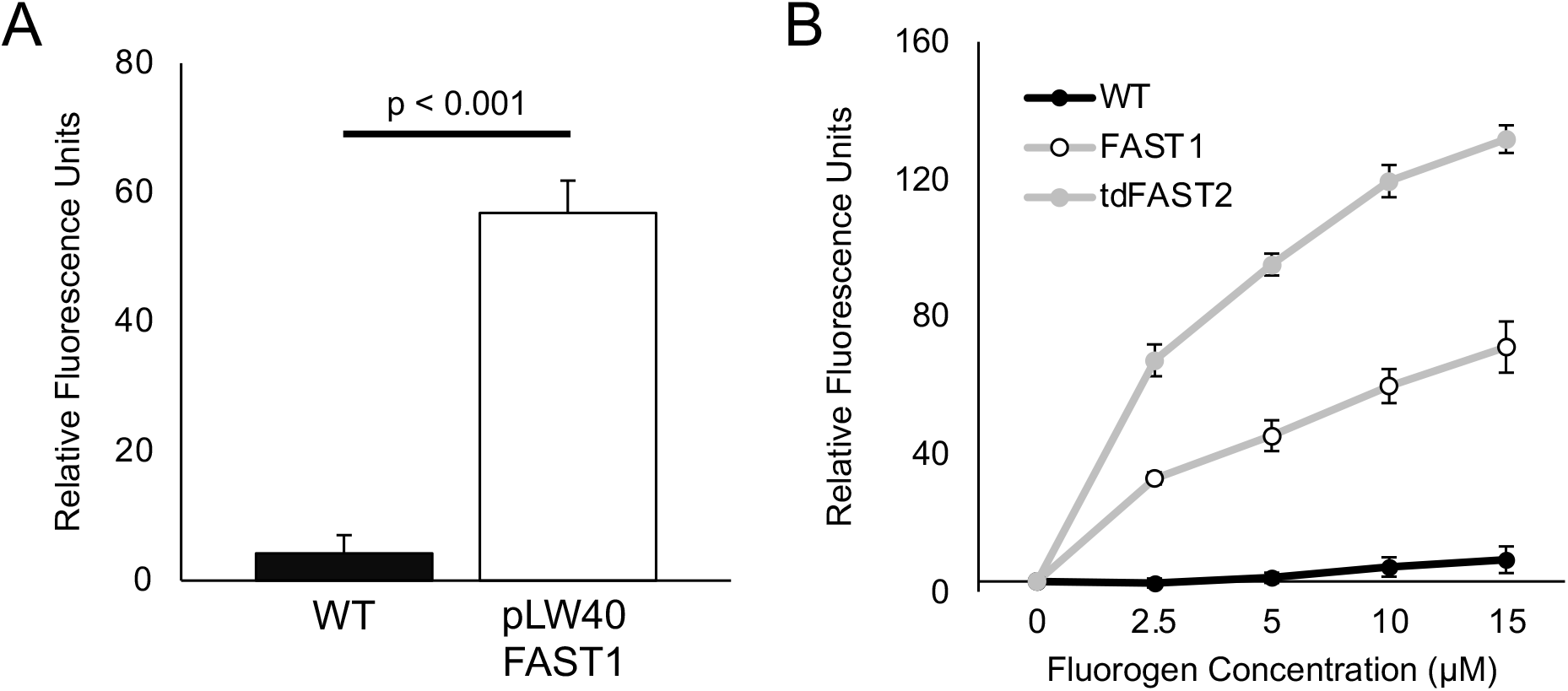
*M. maripaludis* expressing FAST is fluorescent upon HMBR addition **A**. Fluorescence intensities of *M. maripaludis* strains cultivated in McCas medium with H_2_ as the electron donor for growth. HMBR was added to a final concentration of 10 μM. **B**. Titration of HMBR in cells expressing FAST1 or the tandem variant tdFAST2 grown in McCas medium with H_2_. For both panels, relative fluorescence units were determined by normalizing emission readings from a microplate reader against baseline autofluorescence of the sample without fluorogen. Values were also normalized to the OD_600_ of the culture. Data are averages and standard deviations of triplicate measurements.

To further optimize FAST1, we assessed autofluorescence of wild type cells during different stages of growth. We found that cells exhibited the lowest levels of autofluorescence prior to reaching OD_600_ = ∼0.40 or ∼1.0 in medium supplemented with formate or H_2_ as the electron donor for growth, respectively (Fig. S1). We hypothesize that increased autofluorescence at higher OD_600_ is due to an accumulation of oxidized F_420_ when electron donors become growth limiting (17). We also noted that autofluorescence was generally higher in cultures grown on H_2_ and lower in cultures grown on formate; therefore, formate grown cells were used in subsequent experiments when possible. Finally, we tested whether altering concentrations of HMBR could further optimize fluorescence over background. In general, increasing concentrations of HMBR led to increased fluorescence (Fig. 1B). For all subsequent experiments, a concentration of 10 µM HMBR was used unless otherwise indicated.

### Anoxic microscopy with a microscope housed inside of an anoxic chamber

Due to the oxygen sensitivity of *M. maripaludis*, live-cell imaging requires strict anoxia. To overcome this limitation, we developed a platform to perform fluorescence microscopy using an ECHO Revolve R4 hybrid microscope inside of a Coy anaerobic chamber (Fig. 2A). The microscope utilizes a computer tablet camera in place of an eyepiece, and all manipulations can be performed using a touch screen with a capacitive stylus. Using this system, HMBR addition, culture mounting, and imaging can all be performed without introducing oxygen. The microscope was operated in the upright orientation for all experiments.

**Figure 2.**
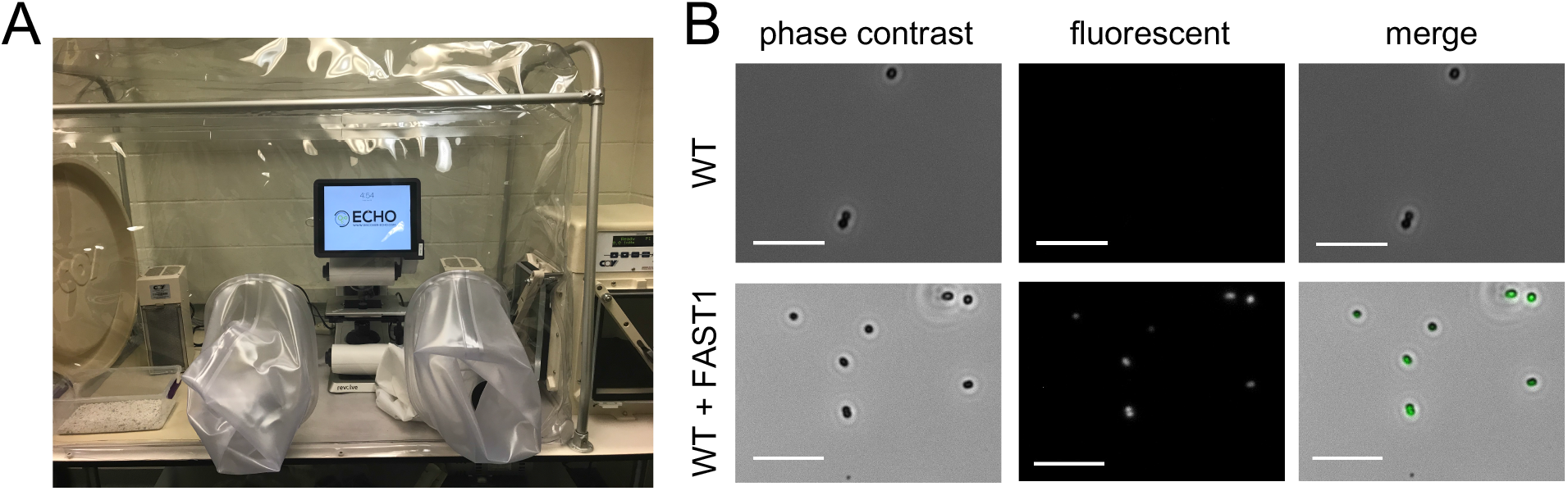
Anoxic microscopy of *M. maripaludis* expressing FAST1. **A**. The anoxic fluorescence microscope used in all experiments. **B**. Images of wild type *M. maripaludis* and a strain expressing FAST1 from the replicating vector pLW40. Scale bars = 10 μm.

Wild type *M. maripaludis* and the strain expressing FAST1 were examined both with and without fluorogen treatment (Fig. 2B). Only FAST1 expressing cells showed noticeable fluorescence gain upon HMBR addition. Because preparing samples in the anoxic chamber involves transferring cells from the high H_2_ (or formate) concentrations of a culture tube to the low hydrogen atmosphere of the chamber (3%), cells were imaged immediately after preparation to mitigate the possibility of increasing autofluorescence from F_420_ oxidation.

### Evaluation of FruA localization and abundance during growth with H_2_ or formate

The F_420_-reducing hydrogenase (Fru) catalyzes the reversible reduction of coenzyme F_420_ to F_420_H_2_ using H_2_ as an electron donor and is the primary source of F_420_H_2_ for methanogenesis (18, 19). The large subunit of the hydrogenase, FruA, was selected as a test case for analyzing a FAST1 translational fusion. Fru is abundant in the cell, shows differential expression with increased abundance when formate is supplied as an electron donor for growth, and is thought to associate with the cell membrane (19–21). Using allelic replacement mutagenesis (1), two strains were created with FAST1 translationally fused to either the N terminus or C terminus of FruA. Both strains were analyzed during early exponential growth with either H_2_ or formate as the electron donor. In general, cells grown in medium with H_2_ exhibited greater autofluorescence when observed under the microscope, and this was true for both N-terminal (FAST1-FruA) and C-terminal (FruA-FAST1) fusions, consistent with the autofluorescence pattern of wild type cells.

The FAST1-FruA construct displayed a pattern of fluorescence consistent with the known membrane localization of Fru (19–21); however, this differed significantly from the pattern observed for the FruA-FAST1 fusion construct (Fig. 3A). Fluorescence in the C-terminal fusion was uniform across the cell while the N-terminal fusion exhibited distinctly higher fluorescence along the outer perimeter of the cell. Distinct foci were regularly observed in the N-terminal fusion strain. The different fluorescence pattern observed between the two FruA constructs is likely due to proteolytic cleavage of the nascent peptide, which is required for maturation of [Ni-Fe] hydrogenases (22). During maturation, Fru is likely proteolytically cleaved at the C terminus, resulting in the loss of the FAST1 tag from FruA-FAST1 fusions and retention of the tag in FAST1-FruA fusions (Fig. 3B). This cleaved FruA-FAST1 expressing strain likely retains FAST1 in the cytoplasm, leading to uniform cellular fluorescence, while the mature, processed, and nonfluorescent hydrogenase localizes to the membrane.

**Figure 3.**
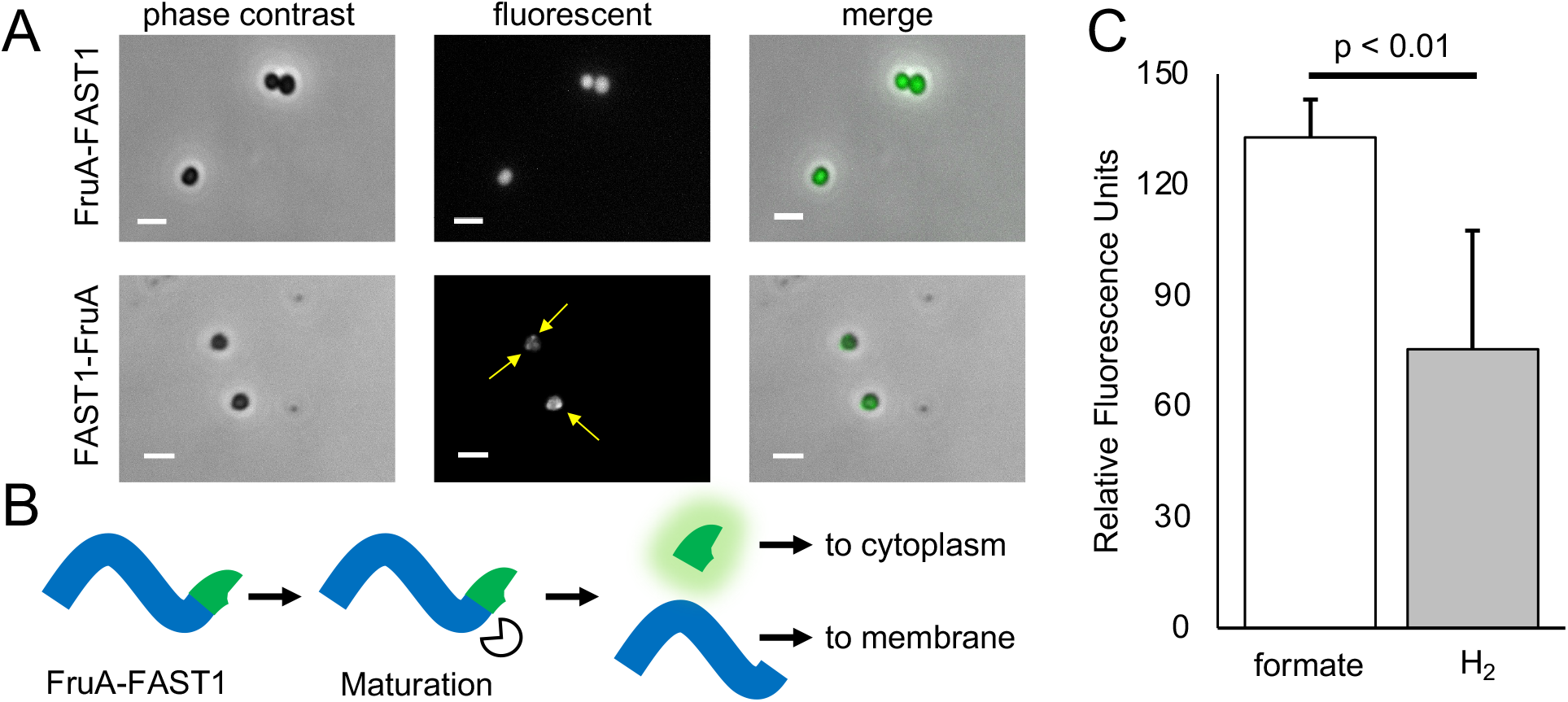
Cellular localization and expression N- and C-terminal translational fusions of FAST1 to FruA **A**. Images of N- and C-terminal FAST fusions to FruA. Scalebars = 2 μm. Arrows indicate locations of fluorescent foci associated with the cell membrane. **B**. Hypothesized maturation of the FruA peptide in the stain expressing *fruA*-FAST1. Maturation of FruA cleaves FAST off of the mature hydrogenase. **C**. Fluorescence intensities of FAST1-FruA cells grown in medium with either formate H_2_ as the sole electron donor. Values were obtained by averaging fluorescence intensities of single cells from each sample via microscopy. Relative fluorescence was normalized to correct for autofluorescence in absence of fluorogen. Data are averages and standard deviations from five independent cultures.

The FAST1-FruA fusion strain was further analyzed for differences in protein abundance between cells grown in medium containing H_2_ or formate. Several transcriptomic and proteomic studies have suggested that Fru is more abundant in cultures grown under conditions where H_2_ concentrations limit growth or when formate is provided as the sole electron donor (23–26). The FAST1-FruA fusion strain was examined by anoxic fluorescence microscopy during early exponential growth (OD_600_ = 0.2 – 0.4). As before, cultures were processed in the absence of oxygen, treated with final concentrations of 10 μM fluorogen, and placed on a glass slide for immediate viewing. To control for autofluorescence, light intensity was measured on a per cell basis in the absence of fluorogen and mean intensities of cells after fluorogen addition were normalized to the autofluorescence baseline. Cells grown utilizing formate as the sole electron donor had 1.76-fold higher fluorescence when compared to cells grown with H_2_ (Fig. 3C), consistent with increased levels of Fru protein when H_2_ concentrations are low.

### Quantifying expression of *fdhAB* with splitFAST and BiFC

Fdh is a multisubunit protein that is essential for growth on formate (27, 28); therefore, we selected Fdh for further validation of FAST1 in measurements of protein abundance. Additionally, as FAST1 is amenable to analysis using BiFC to study protein-protein interactions (13), we generated FdhAB fusion constructs containing split versions of the FAST1 protein. FAST1 can be expressed as two nonfunctional pieces, NFAST (composed of the N terminal 114 amino acids of FAST1) and CFAST (composed of the subsequent 10 amino acids). With splitFAST, two interacting proteins can be tagged with NFAST and CFAST, and, if they colocalize, the pieces will reconstitute in the cell to form a fully functioning protein and fluoresce upon HMBR addition. *M. maripaludis* contains two isoforms of Fdh (29). Fdh1 was selected for analysis because strains lacking *fdh1* are incapable of growth with formate (27, 28) and transcriptomic and proteomic studies suggest that, like Fru, Fdh1 is more abundant in cultures grown under conditions where H_2_ concentrations limit growth or when formate is provided as the sole electron donor (23–26).

An *M. maripaludis* strain with translational fusions encoding FdhA-NFAST and FdhB-CFAST was generated by allelic replacement of the endogenous genes. We additionally generated control strains containing either FdhA1-NFAST and Mtd-CFAST (Mtd is the F_420_-dependent methylenetetrahydromethanopterin dehydrogenase) or FdhB1-CFAST and Mtd-NFAST. Mtd was chosen as a negative control for these studies as Mtd and Fdh are not known to interact and the genes encoding these two proteins exhibit similar patterns of expression (23, 26). Strains containing FdhA1-NFAST and FdhB1-CFAST, FdhA1-NFAST and Mtd-CFAST, or Mtd-NFAST and FdhB1-CFAST were analyzed by anoxic fluorescence microscopy. Relative fluorescence was measured on a per cell basis. Average fluorescence intensity was significantly higher in the strain containing FdhA1-NFAST and FdhB1-CFAST, and both control strains with Mtd fusion constructs displayed similar levels of background fluorescence (Fig. 4A and B).

**Figure 4.**
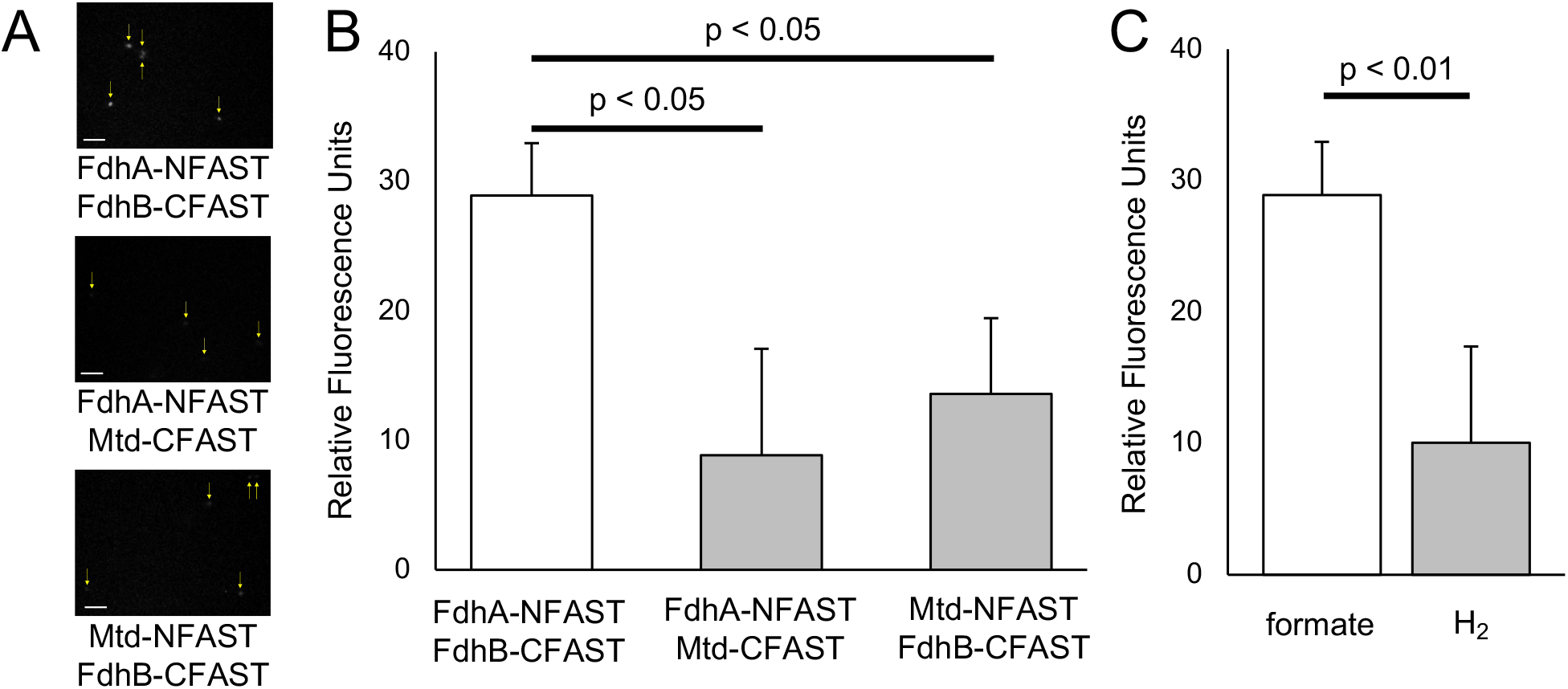
BiFC and fluorescence of Fdh1 with splitFAST **A**. Images splitFAST strains of *M. maripaludis*. Yellow arrows indicate the location of a single cells. Scale bars = 5 μm. **B**. Fluorescence intensities of cells grown expressing various splitFAST constructs. Data were obtained using microscopy and are means and standard deviation of samples collected at least triplicate. **C**. Fluorescence intensities of *fdhA*-NFAST + *fdhB*-CFAST expressing cells grown in medium either formate or H_2_ as the sole electron donor. Data were obtained using microscopy and are means and standard deviation of quadruplicate samples.

To further validate the use of FAST to measure relative protein abundance across growth conditions, the FdhA1-NFAST and FdhB1-CFAST containing strain was analyzed for fluorescence differences between H_2_ and formate grown cells. In cells grown with formate, fluorescence intensity was 2.87-fold higher, consistent with increased Fdh abundance when H_2_ concentrations are low (24) (Fig. 4C).

### Use of tandem FAST2 (tdFAST2) to observe cellular localization of FlaI

In an attempt to further validate FAST for protein localization, we targeted the archaellum (archaeal flagellum) which displays polar localization in intact cells (30). In *M. maripaludis* the major membrane associated components of the archaellum are the anchor (FlaJ) and its associated ATPase (FlaI). Initial attempts to visualize archaella using FAST1 translational fusions were unsuccessful, and we hypothesized that this was due to low fluorescence intensity. To address this, we incorporated a modified version of FAST that has a lower dissociation constant for the fluorogen (FAST2) as a tandem reporter (tdFAST2) (31). This alternative reporter has been shown to achieve higher fluorescence in both bacterial and eukaryotic cells (8, 31).

To test fluorescence, a codon optimized tdFAST2 was transformed into *M. maripaludis* under control of the *M. voltae* histone promoter on the replicating plasmid pLW40neo. FAST1, tdFAST2, and WT were grown to OD_600_ of 0.9, and fluorescence was analyzed using a microplate reader and black, flat bottom 96 well plates across several concentrations of HMBR (Fig. 1B). Strains expressing tdFAST2 displayed significant increases in fluorescence over cells expressing FAST1. When normalized to OD_600_ and controlling for the inherent fluorescence background, microplate reader assays showed that cells expressing tdFAST2 exhibited a 1.9 - 2.1-fold increase fluorescence over FAST1 upon HMBR addition.

A strain was generated expressing FlaI translationally fused to tdFAST2. These cells were viewed by fluorescence microscopy in an anoxic chamber as previously described except that they were treated with 20 μM HMBR. Robust fluorescence was observed associated with the cell membrane and localized to a single focus, consistent with polar localization of archaella (Fig. 5).

**Figure 5.**
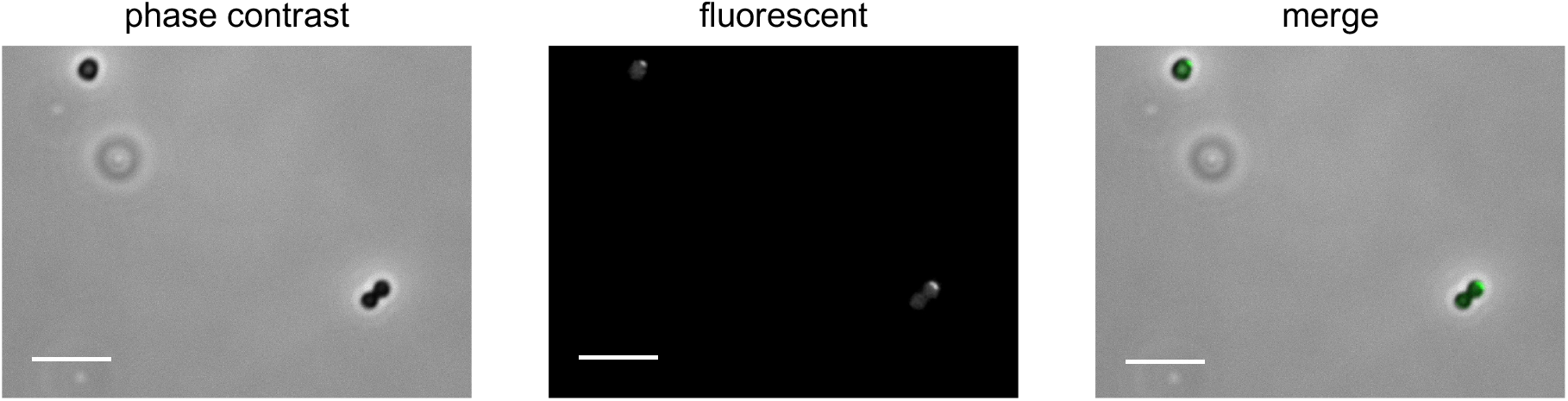
Localization of FlaI tagged with tdFAST2. Images are of early exponential phase cells treated with 20 μM HMBR. Scale bars = 5 μm.

## Discussion

Several features of FAST proteins make them ideal for studies in anaerobic organisms. These include the minimal manipulation required to achieve robust fluorescence, a suite of fluorogens with excitation/emission maxima across the visible spectrum (10), and several variant proteins (e.g. FAST1, FAST2, tdFAST, splitFAST, etc…) to facilitate a variety of studies (11, 13, 31). Additionally, we found that FAST can be used to assess differential protein abundance across growth conditions, protein localization, and protein-protein interactions. Used in conjunction with fluorescence microscopy under anoxic conditions, FAST enables tracking of protein dynamics in live cells of *M. maripaludis*.

FAST proteins function with a variety of fluorogenic ligands. For example, binding of the HBR derivative 4-hydroxy-3,5-dimethoxybenzylidene rhodanine (HBR-3,5DOM) shifts the properties of FAST1 such that ex/em wavelengths are 518/600 nm (32). While HBR-3,5DOM resulted in functional fluorescent protein in *M. maripaludis*, overall fluorescence was lower relative to that of HMBR (Fig. S2), and we generally observed higher background fluorescence from the culture medium, so HBR-3,5DOM was not used further. Autofluorescence was also an issue with cultures grown on H_2_ containing medium, which limited sample collection to exponentially growing cells or cells grown on formate. Autofluorescence was likely due to H_2_ limitation as H_2_ diffusion is outpaced by cellular consumption at higher density; under H_2_ limitation, levels of oxidized, fluorescent F_420_ significantly increase (17). Additionally, removal of cultures from a shaking incubator during sample preparation and transfer to the anoxic chamber can impact H_2_ diffusion, requiring rapid sample preparation and processing. Autofluorescence was less of an issue in cultures grown with formate, likely because formate is soluble in aqueous medium so diffusion does not limit growth.

FAST could be further optimized for use in *M. maripaludis* with the use of protein variants. A previous study with FAST2 and tdFAST2 in mammalian cell culture showed that these variants could achieve 1.7-fold and 3.8-fold higher fluorescence, respectively, compared to the original FAST1 (31). We found a more modest increase for tdFAST2 in *M. maripaludis* of ∼2-fold. FAST2 has identical quantum yield, a very similar molar absorption value, and a lower dissociation constant than FAST1 (31); therefore, it is likely increased fluorescence with tdFAST2 was a result of the tandem nature of the reporter. The performance of tdFAST2 in *M. maripaludis* was consistent with observations in *Eubacterium limosa* where a 1.5-fold increase in brightness was observed (8).

Expression of FAST proteins in *M. maripaludis* could be accomplished using a high expression vector or under control of native promoters on the genome. Generally, total fluorescence was lower when FAST was expressed from native promoters, and cellular fluorescence varied with the use of different promoters. As a result, expression analysis was carried out on single cells using a microscope. For each fused gene, differences in fluorescence were reflective of observed difference in mRNA and protein abundance in previous transcriptomic and proteomic studies (23–26). For Fru, previous studies found that proteins were ∼1.7-fold more abundant when formate was provided as the sole electron donor for growth or when H_2_ concentrations were growth limiting. The FAST-based fluorescence pattern observed here is consistent with these data; mean fluorescence intensities for Fru were 1.76-fold higher when formate was provided as an electron donor. For Fdh, previous studies suggested that these proteins were 2-to 3-fold more abundant when cultures were grown on formate (33), consistent with the 2.87-fold increase in fluorescence for formate grown cells observed here. Together, these data validate the use of FAST protein fusions to measure relative differences in gene expression and protein abundance.

Protein localization applications typically require high fluorescence signal to spatially resolve cellular features. While autofluorescence can complicate these analyses, we successfully visualized the subcellular localization of two proteins that were hypothesized to associate with the cell membrane. Archaella in *M. maripaludis* are associated with the cell pole, and tagging FlaI with tdFAST2 resulted in a single discrete focus characteristic of the hypothesized localization pattern. Localization of Fru to the cell membrane was previously inferred from immunogold labeling of this protein in *M. voltae* (20) and from biochemical studies where Fru activity was enriched in membrane fractions (19). Here we provide a third line of evidence for membrane association of Fru. It remains unclear whether membrane association is required for Fru activity *in vivo*.

Protein fluorescence can be used to characterize protein-protein interactions using dual reporter constructs via fluorescence resonance energy transfer or using a single reporter via BiFC. We used splitFAST to tag both subunits of Fdh and observed robust fluorescence above background through BiFC. While modest increases of fluorescence above background were also observed in control strains expressing either NFAST or CFAST fused to Mtd, fluorescence from NFAST and CFAST fused to FdhA and FdhB was significantly higher than either control experiment. High background in control strains was likely due to the fact that Mtd is most abundant when cultures are grown with formate (23, 24), and BiFC can occur through transient interactions between abundant proteins.

We have demonstrated the utility of FAST for protein analysis in *M. maripaludis*. Combined with anoxic microscopy, FAST allows for live cell imaging, protein localization, determining protein-protein interactions, and expression analysis. These tools should be broadly applicable in other methanogenic archaea with established methods for heterologous protein expression and allelic replacement. Alternative fluorogens and FAST reporter proteins may further expand the utility of FAST in these organisms. The application of anoxic fluorescence reporters presented here expands the already robust toolkit for molecular biology studies in the methanogenic archaea.

## Materials and Methods

### Strains, medium, and growth conditions

Strains in this study are listed in Table S1 in the supplemental material. *M. maripaludis* was grown on McCas medium at 37°C with agitation, except that sodium sulfide was replaced by ammonium sulfide for liquid medium (34, 35). For growth in liquid medium, cultures were grown in 5 ml volumes in Balch tubes. When H_2_ was supplied as the electron donor for growth, Balch tubes were pressurized to 280 kPa with an 80% H_2_, balance CO_2_ gas mix. When formate was supplied as the sole electron donor for growth, McCas-formate medium was used at 37°C without agitation (36) and Balch tubes were pressurized to 210 kPa with an 80% N_2_, balance CO_2_ gas mix. For growth on agar plates, 1.5% noble agar was used and cultures were grown in an anaerobic incubation vessel pressurized to 140 kPa with an 80% H_2_, balance CO_2_ gas mix as described (34). When necessary, the following antibiotics were added to medium at the noted concentrations: neomycin (1 mg ml^−1^), puromycin (2.5 μg ml^−1^), or 6-azauracil (0.25 mg ml^−1^).

### Plasmid construction and transformation of *Escherichia coli*

Plasmids and primers are listed in Table S1 in the supplemental material. To generate FAST constructs, a codon optimized version of the gene was synthesized by Integrated DNA technologies. The codon optimized sequences were determined using the JCat codon optimizer (14) with default parameters with *M. maripaludis* selected as the organism. To generate the tdFAST2 construct, it was necessary to alter the sequence of the gene to prevent loss of the tandem sequence through homologous recombination. To accomplish this, the N-terminal copy of FAST2 was codon optimized in JCat using *M. maripaludis* as the selected organism and the C-terminal copy was codon optimized in JCat using *Methanocaldococcus jannaschii*. The nucleotide sequences of the FAST1 and tdFAST2 genes used in this study can be found in table S2. To further prevent homologous recombination within the tdFAST2 sequence, the 21 base pairs that separate the N-terminal and C-terminal copies were manually edited to reduce similarity to the 33 base pair linker that was consistently used for tagged constructs. In this case, the second most prevalent codon was chosen as determined by the Kazusa database (37).

To generate in-frame FAST constructs, genomic regions flanking either the N terminus or the C terminus of the gene of interest were amplified by PCR using primers suitable for downstream assembly using Gibson assembly (38). Codon optimized FAST1 or tdFAST2 with an additional 33-base pair linker sequence encoding a linker peptide (7) was also amplified by PCR. PCR products were assembled into XbaI/NotI digested pCRuptneo (36). pCRuptneo contains features for propagation in *E. coli* (origin of replication and ampicillin resistance gene) and for selection (neomycin selection) and counterselection (uracil phosphoribosyltransferase) in methanogens. For expression on pLW40 (15), FAST1 or tdFAST2 were PCR amplified and placed into the vector by Gibson assembly at the NsiI and AscI restriction sites under control of the *M. voltae* histone promoter.

Gibson assembled constructs were transferred to *E. coli* DH5α by electroporation. *E. coli* transformants were selected on lysogeny broth agar medium containing ampicillin (50 μg ml^−1^). Plasmids were extracted using the PureLink Quick Plasmid Miniprep kit (Invitrogen) before transfer to *M. maripaludis*. All constructs were sequence verified by Sanger sequencing at the University of Minnesota Genomics Center.

### Transformation of *M. maripaludis*

All strains used in this study were generated in an *M. maripaludis* background lacking the gene for uracil phosphoribosyltransferase (Δ*upt* mutant) (34). DNA was introduced into *M. maripaludis* using either a natural transformation method or a polyethylene glycol (PEG) mediated transformation method (1, 39). For splitFAST constructs, compound, sequential transformations were performed using the integrative vector pCRuptneo. NFAST and CFAST sequences were assembled by amplifying a truncated portion of codon optimized FAST1 sequence. Each splitFAST strain was transformed once with either NFAST or CFAST, subjected to selection and counterselection to remove the integrated pCRuptneo vector, then transformed again to incorporate the second tag.

Cultures for natural transformation were grown in McCas medium using a 2% vol/vol inoculum and H_2_ as the electron donor. After overnight (∼18 h) growth to stationary phase (OD_600_, ∼1), cultures were moved into a room temperature Coy anaerobic chamber (3% H_2_, 10% CO_2_ atmosphere). Plasmid DNA that was preequilibrated for 1 hour in the anaerobic chamber was mixed with 0.5 ml fresh McCas medium and added to the culture using a syringe. The culture/plasmid mixture was pressurized to 280 kPa with an 80% H_2_, balance CO_2_ gas mix and incubated at 37°C with agitation for 4 hours. After outgrowth, 0.2 mls of culture material was transferred to medium containing antibiotic (neomycin for pCRuptneo or puromycin for pLW40) to select for transformants. After growth on neomycin, integrative vectors were allowed to resolve by overnight growth without selection. Mutants were isolated by plating onto 6-azauracil containing McCas agar and resulting colonies were screened by PCR.

PEG-mediated transformations followed the protocol of (1, 39) with all steps performed under anoxic conditions. Briefly, cultures were grown to OD_600_ of ∼0.7 before washing in transformation buffer (TB: 50 mM Tris, 350 mM sucrose, 380 mM NaCl, 1 mM MgCl_2_, pH 7.5). Washed cells were resuspended in 0.375 ml of TB and ∼5 µg of plasmid DNA was added before addition of 0.225 ml PEG solution (40% wt/vol PEG 8000 in TB). After a 1 hour incubation, cells were washed twice in McCas medium, pressurized to 280 kPa with an 80% H_2_, balance CO_2_ gas mix and incubated at 37°C with agitation for at least 4 hours. After outgrowth, cultures were treated the same as for the natural transformation method.

### Fluorescence Quantification Using a SpectraMax M2e plate reader

HMBR was purchased from The Twinkle Factory (https://www.the-twinkle-factory.com/) and solubilized in DMSO to a concentration of 2 mM as a stock solution for all experiments. *M. maripaludis* cells were grown in McCas medium with H_2_ to stationary phase (OD_600_ = ∼0.9) before analysis. *M. maripaludis* liquid culture was directly aliquoted wells of a black, flat bottom 96 well plate (Bioassay systems #P96FL), with 10 μM of HMBR (unless otherwise indicated in the text) to a final volume of 200 μl. Cells were incubated with HMBR for 60 seconds per the manufacturer’s instructions before readings. Fluorescence was measured in a SpectraMax M2e plate reader (Molecular Devices) and analyzed in SoftMax Pro 7 software. Excitation wavelength was set to 481 nm and emission wavelength was set to 540 nm utilizing auto emission cutoff settings. Each sample was normalized to the baseline reading of the same sample without fluorogen addition to correct for autofluorescence. Samples were further normalized on a per cell basis using OD_600_ of the cells prior to plate preparation.

Samples prepared with HBR-3,5DOM utilized the same methodology as samples treated with HMBR with the addition of a wash step. To reduce the high levels of autofluorescence in the media, cells were pelleted aerobically via centrifugation at 15,500 x g for two minutes and resuspended to their original volume with sterile ddH_2_O.

### Microscopy

Imaging was performed in a Coy-type anaerobic chamber with an environment of 3% H_2_, balance N_2_. The chamber also contained a 20 cm x 20 cm tray with drierite desiccant to control humidity. All imaging was performed using an ECHO Revolve R4 hybrid microscope operated in the upright orientation with a high resolution condenser (numerical aperture 0.85, working distance 7 mm). Images were taken with the high gain setting on and exposure settings were modified depending on the genetic construct. Cells were viewed using a 40x fluorite objective lens (NA 0.75 and 0.51 mm) or a 100x fluorite oil phase objective lens (NA 1.30 and 0.2 mm). Fluorescence imaging was carried out using standard FITC filter sets.

To minimize autofluorescence, culture tubes were kept in the incubator until they were ready for immediate imaging. Samples were prepared by aliquoting liquid culture into microcentrifuge tubes and subsequently adding HMBR in DMSO to final concentrations as specified. Aliquots of 5 μl were added onto glass slides and glass coverslips were placed over them. All steps were performed under anoxic conditions.

### Quantification of Fluorescence Microscopy Data using ImageJ

For expression analysis, 16 bit .tiff images of both a phase contrast and a fluorescent channel were used to analyze a single field of view utilizing a 40x objective. The FIJI package of ImageJ was used (version 2.1.0/1.53j Java ver. 1.8.0_172). A mask was generated using the phase contrast image by generating an 8-bit image using the threshold tool. The ‘watershed’ and ‘fill holes’ binary features were used as appropriate. The pixels of the resulting image in which cells were located were assigned a value of 1 and pixels elsewhere were assigned a value of 0 through the use of the division function. This image was multiplied with the corresponding fluorescence image to create a composite image with 32-bit float, and the mean fluorescence intensity per cell was determined using the particle analyzer feature. An average of averages was obtained from the compiled list of mean fluorescence intensities. Relative fluorescence units were determined by measuring the gain of fluorescence in cells after HMBR treatment. The average autofluorescent background was determined on a per-sample basis through the methods listed above and was subtracted from the values obtained after the addition of fluorogen. For protein localization analysis, 16-bit .tiff images of both a phase contrast and a fluorescence channel were used to analyze a single field of view utilizing a 100x objective. For figures 3A, 4A, and 5, a median filter was applied for visual clarity.

### Statistical Analysis

Data are presented as means ± standard deviation. Statistical analysis was completed using a two tailed Student *t* test using Microsoft Excel software. P values for Fdh splitFAST against negative controls were determined using a one way ANOVA followed up by a post hoc Dunnett’s test for multiple comparison significance in R (version 3.6.2) (40). Significant differences were considered when *P* values were <0.05.

## Acknowledgements

We thank Thomas Hanson for suggesting FAST as a potential reporter system and providing HMBR for preliminary experiments. This work was sponsored by a Young Investigator Program award from the Army Research Office, grant number W911NF-19-1-0024.

## References

1. Sarmiento FB, Leigh JA, Whitman WB. 2011. Genetic systems for hydrogenotrophic methanogens. Meth Enzymol 494:43–73.

2. Leigh JA, Albers S-V, Atomi H, Allers T. 2011. Model organisms for genetics in the domain Archaea: methanogens, halophiles, Thermococcales and Sulfolobales. FEMS Microbiol Rev 35:577–608.

3. Thauer RK, Kaster A-K, Seedorf H, Buckel W, Hedderich R. 2008. Methanogenic archaea: ecologically relevant differences in energy conservation. Nat Rev Microbiol 6:579–591.

4. Purwantini E, Mukhopadhyay B. 2009. Conversion of NO_2_ to NO by reduced coenzyme F_420_ protects mycobacteria from nitrosative damage. Proc Natl Acad Sci USA 106:6333–6338.

5. Heim R, Prasher DC, Tsien RY. 1994. Wavelength mutations and posttranslational autoxidation of green fluorescent protein. Proc Natl Acad Sci USA 91:12501–12504.

6. Buckley AM, Petersen J, Roe AJ, Douce GR, Christie JM. 2015. LOV-based reporters for fluorescence imaging. Curr Opin Chem Biol 27:39–45.

7. Plamont M-A, Billon-Denis E, Maurin S, Gauron C, Pimenta FM, Specht CG, Shi J, Quérard J, Pan B, Rossignol J, Moncoq K, Morellet N, Volovitch M, Lescop E, Chen Y, Triller A, Vriz S, Le Saux T, Jullien L, Gautier A. 2016. Small fluorescence-activating and absorption-shifting tag for tunable protein imaging in vivo. Proc Natl Acad Sci USA 113:497–502.

8. Flaiz M, Ludwig G, Bengelsdorf FR, Dürre P. 2021. Production of the biocommodities butanol and acetone from methanol with fluorescent FAST-tagged proteins using metabolically engineered strains of Eubacterium limosum. Biotechnol Biofuels 14:117.

9. Streett HE, Kalis KM, Papoutsakis ET. 2019. A strongly fluorescing anaerobic reporter and protein-tagging system for Clostridium organisms based on the fluorescence-activating and absorption-shifting tag protein (FAST). Appl Environ Microbiol 85:e00622–19.

10. Myasnyanko IN, Gavrikov AS, Zaitseva SO, Smirnov AY, Zaitseva ER, Sokolov AI, Malyshevskaya KK, Baleeva NS, Mishin AS, Baranov MS. 2020. Color tuning of fluorogens for FAST fluorogen-activating protein. Chem Eur J. 27:3986–3990.

11. Tebo AG, Moeyaert B, Thauvin M, Carlon-Andres I, Böken D, Volovitch M, Padilla-Parra S, Dedecker P, Vriz S, Gautier A. 2021. Orthogonal fluorescent chemogenetic reporters for multicolor imaging. Nat Chem Biol 17:30–38.

12. Monmeyran A, Thomen P, Jonquière H, Sureau F, Li C, Plamont M-A, Douarche C, Casella J-F, Gautier A, Henry N. 2018. The inducible chemical-genetic fluorescent marker FAST outperforms classical fluorescent proteins in the quantitative reporting of bacterial biofilm dynamics. Sci Rep 8:10336.

13. Tebo AG, Gautier A. 2019. A split fluorescent reporter with rapid and reversible complementation. Nat Commun 10:2822.

14. Grote A, Hiller K, Scheer M, Münch R, Nörtemann B, Hempel DC, Jahn D. 2005. JCat: a novel tool to adapt codon usage of a target gene to its potential expression host. Nucleic Acids Res 33:W526–31.

15. Dodsworth JA, Leigh JA. 2006. Regulation of nitrogenase by 2-oxoglutarate-reversible, direct binding of a PII-like nitrogen sensor protein to dinitrogenase. Proc Natl Acad Sci USA 103:9779–9784.

16. Gardner WL, Whitman WB. 1999. Expression vectors for Methanococcus maripaludis: overexpression of acetohydroxyacid synthase and beta-galactosidase. Genetics 152:1439–1447.

17. de Poorter LMI, Geerts WJ, Keltjens JT. 2005. Hydrogen concentrations in methaneforming cells probed by the ratios of reduced and oxidized coenzyme F_420_. Microbiology 151:1697–1705.

18. Hendrickson EL, Leigh JA. 2008. Roles of coenzyme F_420_-reducing hydrogenases and hydrogen-and F_420_-dependent methylenetetrahydromethanopterin dehydrogenases in reduction of F_420_ and production of hydrogen during methanogenesis. J Bacteriol 190:4818–4821.

19. Baron SF, Ferry JG. 1989. Purification and properties of the membrane-associated coenzyme F_420_-reducing hydrogenase from Methanobacterium formicicum. J Bacteriol 171:3846–3853.

20. Muth E. 1988. Localization of the F_420_-reducing hydrogenase in Methanococcus voltae cells by immuno-gold technique. Arch Microbiol 150:205–207.

21. Baron SF, Williams DS, May HD, Patel PS, Aldrich HC, Ferry JG. 1989. Immunogold localization of coenzyme F_420_-reducing formate dehydrogenase and coenzyme F_420_-reducing hydrogenase in Methanobacterium formicicum. Arch Microbiol 151:307–313.

22. Lacasse MJ, Zamble DB. 2016. [NiFe]-hydrogenase maturation. Biochemistry 55:1689–1701.

23. Costa KC, Yoon SH, Pan M, Burn JA, Baliga NS, Leigh JA. 2013. Effects of H_2_ and formate on growth yield and regulation of methanogenesis in Methanococcus maripaludis. J Bacteriol 195:1456–1462.

24. Xia Q, Wang T, Hendrickson EL, Lie TJ, Hackett M, Leigh JA. 2009. Quantitative proteomics of nutrient limitation in the hydrogenotrophic methanogen Methanococcus maripaludis. BMC Microbiol 9:149.

25. Hendrickson EL, Liu Y, Rosas-Sandoval G, Porat I, Söll D, Whitman WB, Leigh JA. 2008. Global responses of Methanococcus maripaludis to specific nutrient limitations and growth rate. J Bacteriol 190:2198–2205.

26. Hendrickson EL, Haydock AK, Moore BC, Whitman WB, Leigh JA. 2007. Functionally distinct genes regulated by hydrogen limitation and growth rate in methanogenic Archaea. Proc Natl Acad Sci USA 104:8930–8934.

27. Costa KC, Lie TJ, Xia Q, Leigh JA. 2013. VhuD facilitates electron flow from H_2_ or formate to heterodisulfide reductase in Methanococcus maripaludis. J Bacteriol 195:5160–5165.

28. Lupa B, Hendrickson EL, Leigh JA, Whitman WB. 2008. Formate-dependent H_2_ production by the mesophilic methanogen Methanococcus maripaludis. Appl Environ Microbiol 74:6584–6590.

29. Poehlein A, Heym D, Quitzke V, Fersch J, Daniel R, Rother M. 2018. Complete genome sequence of the Methanococcus maripaludis type strain JJ (DSM 2067), a model for selenoprotein synthesis in Archaea. Genome Announc 6:e00237–18.

30. Jarrell KF, Stark M, Nair DB, Chong JPJ. 2011. Flagella and pili are both necessary for efficient attachment of Methanococcus maripaludis to surfaces. FEMS Microbiol Lett 319:44–50.

31. Tebo AG, Pimenta FM, Zhang Y, Gautier A. 2018. Improved chemical-genetic fluorescent markers for live cell microscopy. Biochemistry 57:5648–5653.

32. Li C, Plamont M-A, Sladitschek HL, Rodrigues V, Aujard I, Neveu P, Le Saux T, Jullien L, Gautier A. 2017. Dynamic multicolor protein labeling in living cells. Chem Sci 8:5598–5605.

33. Wood GE, Haydock AK, Leigh JA. 2003. Function and regulation of the formate dehydrogenase genes of the methanogenic archaeon Methanococcus maripaludis. J Bacteriol 185:2548–2554.

34. Fonseca DR, Abdul Halim MF, Holten MP, Costa KC. 2020. Type IV-like pili facilitate transformation in naturally competent archaea. J Bacteriol. 202:e00355–20.

35. Moore BC, Leigh JA. 2005. Markerless mutagenesis in Methanococcus maripaludis demonstrates roles for alanine dehydrogenase, alanine racemase, and alanine permease. J Bacteriol 187:972–979.

36. Costa KC, Wong PM, Wang T, Lie TJ, Dodsworth JA, Swanson I, Burn JA, Hackett M, Leigh JA. 2010. Protein complexing in a methanogen suggests electron bifurcation and electron delivery from formate to heterodisulfide reductase. Proc Natl Acad Sci USA 107:11050–11055.

37. Nakamura Y, Gojobori T, Ikemura T. 2000. Codon usage tabulated from international DNA sequence databases: status for the year 2000. Nucleic Acids Res 28:292.

38. Gibson DG, Young L, Chuang R-Y, Venter JC, Hutchison CA, Smith HO. 2009. Enzymatic assembly of DNA molecules up to several hundred kilobases. Nat Methods 6:343–345.

39. Tumbula DL, Makula RA, Whitman WB. 1994. Transformation of Methanococcus maripaludis and identification of a Pst I-like restriction system. FEMS Microbiol Lett 121:309–314.

40. R Core Development Team. 2019. A language and environment for statistical computing. R Foundation for Statistical Computing, Vienna, Austria. https://www.r-project.org/.

